# Genetic Diversity, Population Structure, and Cannabinoid Variation in Feral *Cannabis sativa* Germplasm from the United States

**DOI:** 10.1101/2025.03.10.642411

**Authors:** Ademola Aina, Jonathan P Wenger, Eliot Stanton, Chandrani Gon Majumdar, Mahmoud ElSohly, George D Weiblen, Shelby Ellison

## Abstract

*Cannabis sativa* is one of the earliest plants to be domesticated for fiber, food and medicine. Seed from *Cannabis* grown for industrial purposes during the 18^th^ through 20^th^ centuries have escaped production and established feralized populations across the United States. To maximize the potential of feral *Cannabis* germplasm, determining the genetic structure and cannabinoid profile is crucial for selection and breeding of new compliant regionally adapted hemp cultivars. To resolve this, a collection of feral *Cannabis*, comprising 760 plants across twelve US states were sequenced using Genotyping-by-Sequencings (GBS), genotyped at the *cannabinoid synthase* (*CBDAS)* gene, and subject to gas chromatography-mass spectrometry (GC-MS) to assess cannabinoid profiles. Clustering analyses by ADMIXTURE and Principal Component Analysis (PCA) stratified the germplasm into five clusters (Mississippi-River, West North Central-b, West North Central-a, New York, and Indiana). The cannabinoid genotyping assay resolved the feral collections into Type I - C^X^C^X^ (6%), Type II - C^F^C^X^ (15%), and Type III - C^F^C^F^ (78%). Total cannabinoid content ranged from 0.21% to 4.73%. The assessment of genetic diversity, population structure, and cannabinoid profile of the US feral *Cannabis* collection provides critical information and germplasm resources to develop new and improve existing hemp cultivars.

## INTRODUCTION

*Cannabis sativa* L. is a versatile crop that has been cultivated for thousands of years for its bast fiber (stem) and hurd fiber (stem pith), edible grain (seed) and cannabinoids (flowers). *Cannabis* plants produce a substantial number of chemical compounds known as cannabinoids. These cannabinoids comprise C21 terpenophenolic compounds responsible for the medicinal and psychotropic effects obtained from *Cannabis* use^1–3^. There are over a hundred cannabinoids^4^ but the two main molecules of interest have always been delta-9-tetrahydrocannabinol (THC) and cannabidiol (CBD), both derived from the same precursor, cannabigerolic acid (CBGA)^5^. Consumption of high THC (drug-type *Cannabis*) results in intoxicating and/or analgesic effects^6^, while hemp-type *Cannabis* contains predominantly non-intoxicating cannabinoids (CBD). A single locus with codominant alleles purportedly described the pattern of inheritance for THC to CBD ratios in *Cannabis* plants^7,8^. Recent findings have resolved mutually exclusive chromosome 7 haplotypes exhibiting orthogonal genetic and physical linkage of functional vs. non-functional *THCAS* and *CBDAS* loci^9–11^. Classification of *Cannabis* as hemp or drug-type, is based on the concentration of THC, with 0.3% THC as the allowable upper threshold of industrial hemp (hemp-type *Cannabis*) as defined by the 2014 and 2018 Farm Bills^12,13^.

Hemp was introduced to North America in the early 17th century^14^, and was used to make ropes, grain bags, Conestoga wagon covers, and clothing. In the early 20^th^ century, a USDA agriculturalist Lyster Dewey, began researching industrial hemp production to help meet the US demand for a domestic fiber source with initial breeding efforts aimed at improving fiber quality and yield^15,16^. During this period, industrial hemp production was primarily concentrated in the Midwest region of the US due to favorable growing conditions and accessible hemp processing mills. Production peaked in 1943 when approximately 176,000 acres were cultivated to supply fiber for the World War II war effort^17^. After the war, all *Cannabis* effectively became impractical to grow because of the resumed enforcement of the 1937 Marihuana Tax Act and production rapidly halted. Despite the cessation of cultivation, feralized industrial hemp populations were established after escaping production and continued reproducing in the wild across the major prior production regions in the US^18^. Geographically, these remnants of feral *Cannabis* populations are widely dispersed and grow in a wide range of climatic conditions. Feral *Cannabis* thrives under full exposure to sunlight and in nitrogen-rich, well-drained soils where its extensive and deep root structure absorbs enough water for the shoots to withstand hot and dry weather^19^. As of 2025, iNaturalist and Global Biodiversity Information Facility (GBIF) online databases currently recorded, 1,795 and 1,207 human observations of feral *Cannabis* in the US, respectively. Many of these observations occurred in the Midwest and Northeast areas of the US where they were previously cultivated in large acreages. Feral hemp is commonly found growing in disturbed, nutrient-rich lands including pastures, feed lots, farmyards, cultivated field borders, open woods, flood plains, dumpsites, roadsides, and railroad rights-of-way.

The recent legalization of hemp, with its many potential industrial and medicinal applications, has reinvigorated the scientific investigation of *Cannabis*. Significant scientific advancements have been made in *Cannabis* genetics and genomics over the last decade. This includes the completion of a several high-quality genome assemblies^9,20,21^, understanding genetic relatedness of major market classes, and studies on the genetic underpinnings of cannabinoid inheritance, sex determination, and many agronomic and morphological traits^22–24^. *Cannabis* research and breeding efforts lag behind that of many modern crops. The urgency of creating and disseminating industrial hemp genetic resources cannot be overemphasized, with these tasks falling on genebanks to ensure adequate germplasm supply. Despite many germplasm repositories across the globe, only a handful of them show records of deliberate conservation of *Cannabis* accessions. Governments and universities have been the custodians of large-scale *ex-situ* germplasm collections for centuries. One such government-owned collection is the USDA-ARS National Plant Germplasm System (NPGS), the world’s largest supplier of germplasm, holding 617,467 accessions representing 17,482 species^25^. As a result of federal prohibition starting in 1937, hemp germplasm has been excluded from such conservation until the 2021 establishment of the USDA-ARS hemp germplasm repository in Geneva, New York. The current collection holds 581 *Cannabis sativa* accessions of which 57 are available for distribution.

Escaped, naturalized, and regionally adapted feral cannabis populations that exist across the US represent a potential source of untapped genetic diversity. These populations potentially harbor a wide range of genetic variation lost in cultivated crops, providing opportunities to introduce new traits for improved crop performance. As crops become feral, traits suited to local conditions like drought, extreme temperatures, poor soil, could be selected for. These adaptations are valuable for developing climate-resilient crops by incorporating genetic material from feral populations via breeding. Moreover, studying feral populations offers insights into the evolutionary processes of crop domestication and artificial selection. Several studies have described the population structure and chemical profiles in diverse *C. sativa* collections^26–32^ by using next-generation sequencing methods on mostly outsourced drug type *Cannabis* from Canada, Europe or Asian origin. Previous studies from four US states (Minnesota, Nebraska, Kansas and Colorado) have explored feral *Cannabis* diversity and found feral plants to be a rich source of genetic diversity^33–35^. The goal of this study is to populate the recently established USDA-ARS hemp germplasm repository in Geneva, NY, with genetically characterized, diverse, and compliant plant material to provide a valuable resource for the scientific community. To complete this goal we executed a large-scale collection effort to collect, genotype, and chemotype unique feral *Cannabis* germplasm from across the US to support research efforts, conserve biodiversity and develop regionally adapted, compliant hemp cultivars.

## RESULTS

### Sample collection

Seeds and floral tissues from 1,821 feral plants were collected across 12 states and 91 unique populations, in the late summer or early fall of 2022 and 2023 (Figure 1a and 1b; Supplemental Table 1). The largest collection of samples came from Wisconsin (n = 589) and the smallest from Colorado (n = 11).

**Figure 1.**
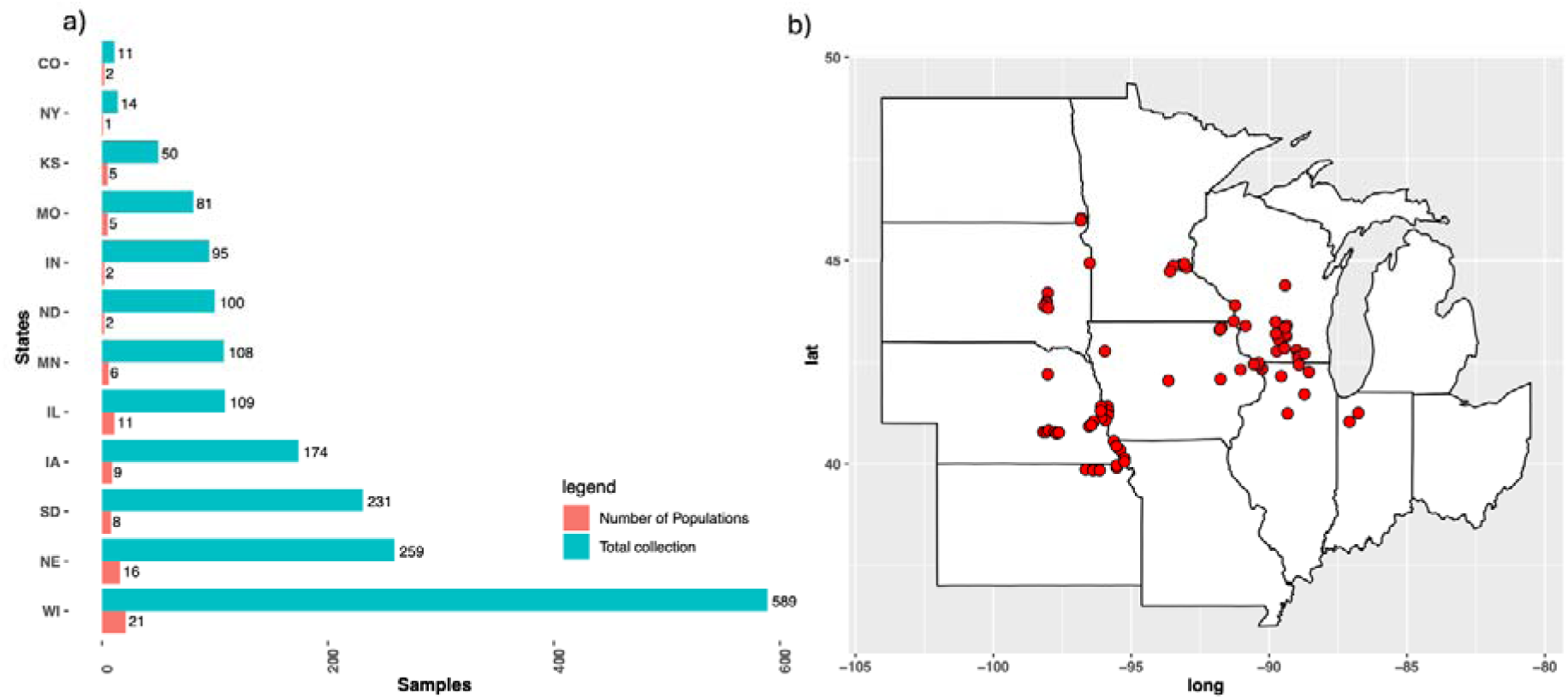
Feral *Cannabis* collection. (a) Bar plot showing total number of individual feral *Cannabis* accessions collected by state alongside the number of populations sampled. (b) Midwestern map showing geographical distribution of sampled locations (Colorado and New York excluded).

### SNP distribution

Among the 91 sampled populations, 5-20 individual plants were selected for Genotyping-By-Sequencing (GBS). A total of 2,739,241,114 barcoded reads were aligned with the *Cannabis* reference genome^9^ resulting in identification of 159,382 polymorphic SNPs. After filtering, the final dataset included 21,037 high quality SNPs. SNP counts ranged from 1,386 on chromosome 7 to 2,829 on chromosome 1 (Supplemental Table 2), with well distributed coverage across all chromosomes (Supplementary Figure 1).

### Population Structure

The cross-validation method by ADMIXTURE was unable to resolve the optimum number of clusters (Supplemental Figure 2) for the samples. However, the Bayesian Information Criterion (BIC) value indicated a rapid decline until K = 5 clusters, after which the value began to rise, suggesting that five groups should be retained (Figure 2a). Following ADMIXTURE analysis, individual feral *Cannabis* accessions were assigned into a group based on their maximum proportion of membership (Q) value (Figure 2b). To avoid groups with recent admixture between populations or cultivated *Cannabis*, a > 70 percent membership threshold was used to assign subpopulations identity, retaining 346 samples (Supplementary Table 3). Group 1 (Mississippi-River) had the largest number of samples with a total of 177 accessions largely composed of feral collections from eight states (MN, WI, IL, MO, SD, IA, NE and KS) and dominated by accessions from WI (57), MN (52), and IL (36) (Supplementary Table 4). Group 2 (West North Central-a) comprised 96 accessions from SD, NE, IA, ND, CO, KS, MO, IL and MN, however SD (40 accessions), NE (17 accessions), and ND (12 accessions) were the most represented in this group. Group 3 (New York) comprised 14 samples from New York. Group 4 (West North Central-b) comprised 37 samples, with accessions from SD (11), IA (8) and NE (7), similar to Group 2. Finally, group 5 (Indiana) comprised 22 accessions from Indiana. Principal component analysis (PCA) of the ∼21K SNPs sorted the feral *Cannabis* accessions into the same five major groups (Figure 3a). The majority of the variation explained came from New York and Indiana, respectively, while the other groups clustered tightly. Samples mostly grouped by states with a few minor exceptions (Figure 3b and 3c).

**Figure 2.**
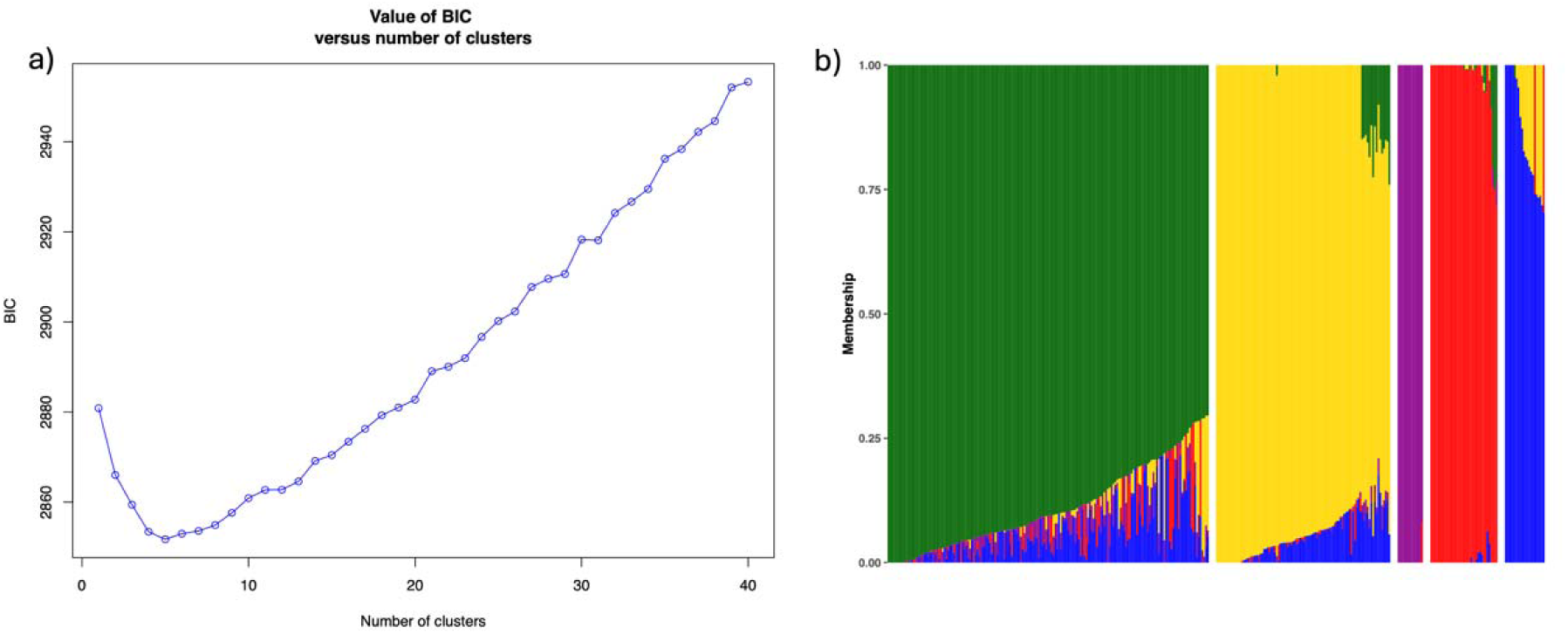
Population ADMIXTURE. (a) Graph of Bayesian Information Criterion versus number of clusters. (b) Admixture ancestry of 346 feral *Cannabis* accessions representing Group 1 (Mississippi-River), Group 2 (West North Central-a), Group 3 (New York), Group 4 (West North Central-b), and Group 5 (Indiana), respectively (from left to right).

**Figure 3.**
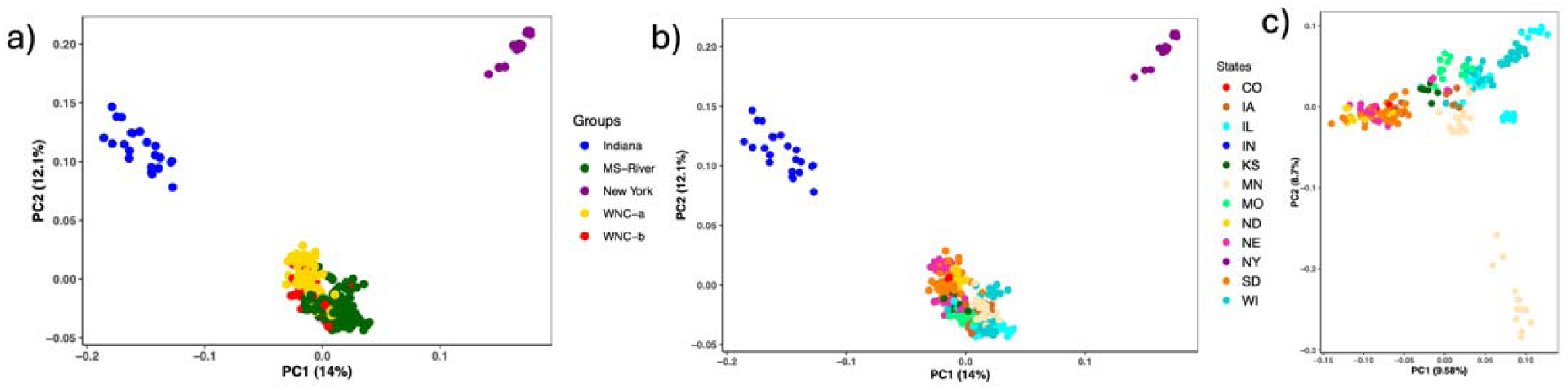
Principle Component Analysis of (a) 346 feral *Cannabis* plants colored by ADMIXTURE clusters, (b) collection states, and (c) collection states after removing Indiana and New York samples.

### Genetic Differentiation

The F_ST_ values between the inferred groups ranged from 0.07 to 0.39 (Table 1). Pairwise genetic differentiation was highest between the New York and Indiana groups (F_ST_ = 0.39), with very low values (F_ST_ = 0.07-0.09) between Mississippi-River, West North Central-a and West North Central-b. Sliding window F_ST_ analysis revealed extremely high F_ST_ values on chromosome 4 and the X chromosome between groups (Figure 4a and 4b, Supplementary Figure 3). The SNPs associated with elevated F_ST_ (65 – 90 Mb) were analyzed separately by PCA showing three discrete clusters (Supplementary Figure 4a and 4b) explaining 63.9% of the observed variation. Inspection of this region indicated a large genome rearrangement with respect to the Cs10 genome that has been observed previously in feral samples^21,34^. The inversion genotype (+/+) was characterized for each of the five groups and was found at 93% and 91% in the West North Central-a (Group 2) and Indiana (Group 5) populations, respectively (Table 2).

**Figure 4.**
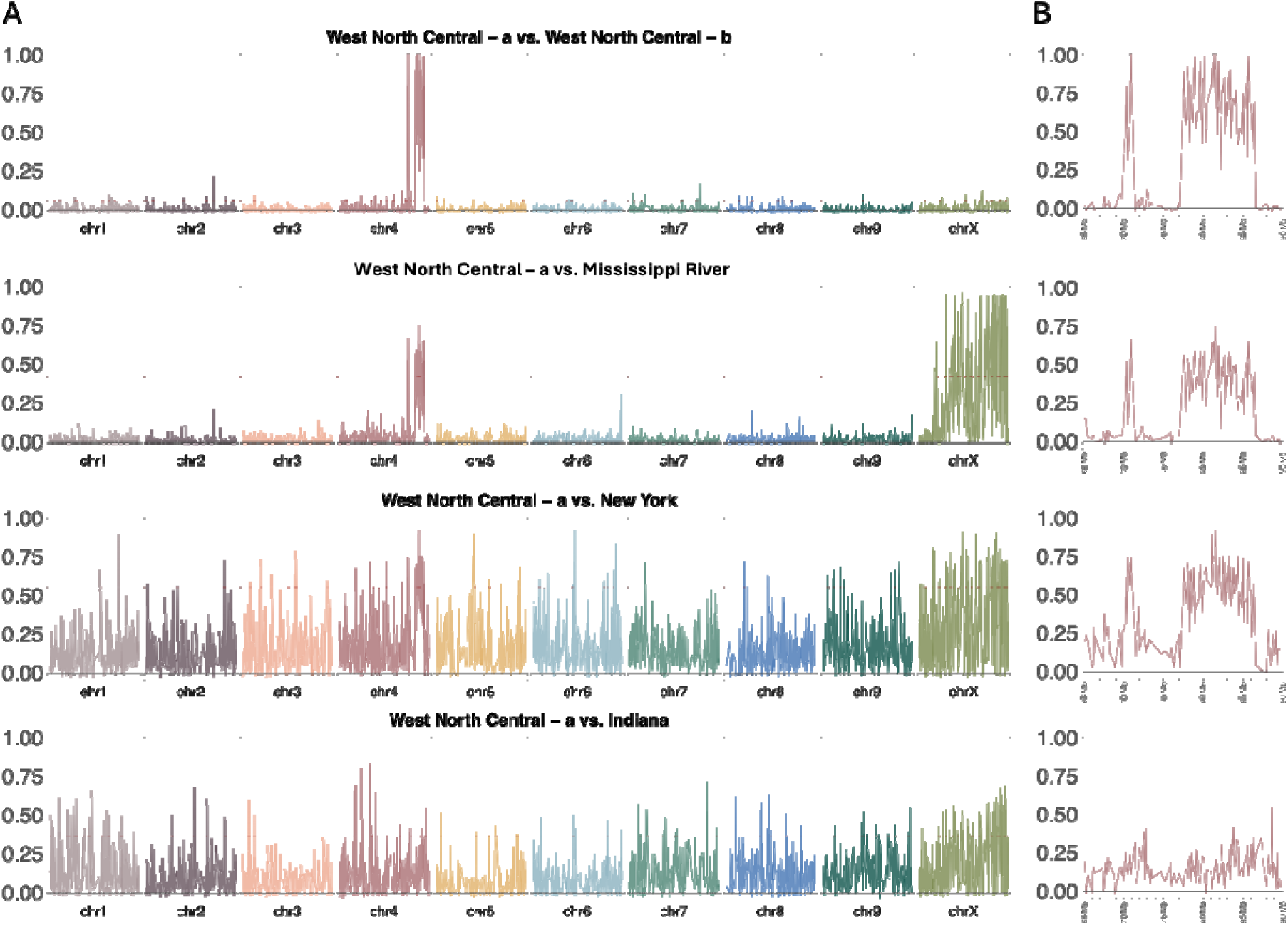
Genetic differentiation between all groups as compared to the “West North Central - a” group as determined by sliding window F_ST_ analysis of 100 kb bins across a) the entire genome and b) a highly differentiated region (65-90 Mb) on Chromosome 4. The red dashed lines indicates the top 5% of F_ST_ values for each comparision.

**Table 1.** Pairwise comparison of genetic differentiation (F_ST_ values) among the five observed groups.

|  | MS-River | WNC-a | New York | WNC-b |
| --- | --- | --- | --- | --- |
| WNC-a | 0.09 |  |  |  |
| New York | 0.22 | 0.25 |  |  |
| WNC-b | 0.07 | 0.07 | 0.28 |  |
| Indiana | 0.17 | 0.14 | 0.39 | 0.20 |

\*Mississippi-River (MS-River), West North Central-a, West North Central-b

**Table 2.**
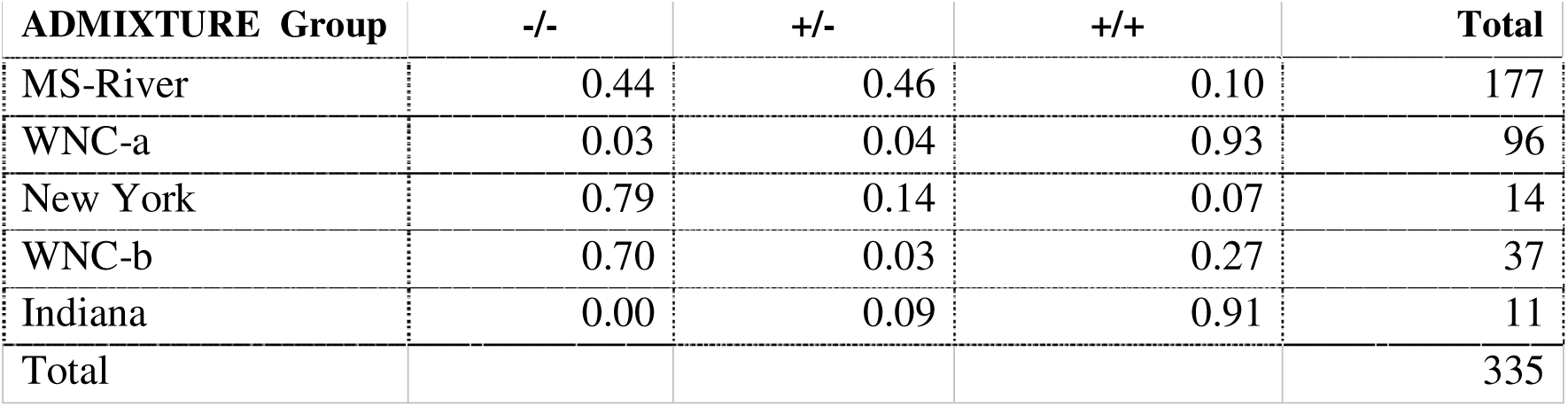
Frequency of chromosome 4 inversion across subpopulations. -/- indicates same as Cs10 reference (no inversion), -/+ is heterozygous for the inversion and +/+ is homozygous for the inversion.

### Genetic Diversity Analysis

Minor allele frequency (MAF) observed and expected heterozygosity (*Ho* and *He*), polymorphism information content (PIC), and the inbreeding coefficient (F_IS_) are used as indicators of the genetic diversity within a population^36^. The average *Ho*, *He*, MAF, PIC, F_IS_ of the 346 accessions were 0.20, 0.20, 0.19, 0.26 and −0.03, respectively (Table 3). Analysis of the genetic diversity of the five *Cannabis* groups showed MAF ranged between 0.12 - 0.22, while polymorphic information content (PIC) was between 0.15 - 0.30. The *Ho*, MAF and PIC of the New York population were lower than those of the Midwestern populations. Hierarchical Analysis of Molecular Variance (AMOVA) based on ADMIXTURE groupings found approximately 9% variation between groups, 20% between accessions that were clustered within groups and extensive variation (71%) among all samples (Supplementary Table 5).

**Table 3.** Diversity indices for the entire feral *Cannabis* collection and by subpopulation. *Ho* = Observed heterozygosity, *He* = Expected heterozygosity, MAF = Minor allele frequency, PIC = Polymorphic information content, F_IS_ = Inbreeding coefficient.

|  | *Ho* | *He* | MAF | PIC | *F<sub>IS</sub>* |
| --- | --- | --- | --- | --- | --- |
| MS-River | 0.24 | 0.24 | 0.22 | 0.30 | -0.08 |
| WNC-a | 0.24 | 0.24 | 0.22 | 0.30 | -0.05 |
| New York | 0.09 | 0.10 | 0.12 | 0.15 | 0.06 |
| WNC-b | 0.24 | 0.23 | 0.22 | 0.30 | -0.07 |
| Indiana | 0.19 | 0.20 | 0.18 | 0.25 | 0.00 |
| All samples | 0.20 | 0.20 | 0.19 | 0.26 | -0.03 |

**Table 4.** Cannabinoid content and genotype frequencies among feral *Cannabis* populations. Genotype frequencies are reported for the *CBDAS* locus corresponding to the three major cannabinoid ratio classes, CBD-type (C^F^C^F^), intermediate (C^F^C^X^), and THC-type (C^X^C^X^). Cannabinoid content, including total cannabinoid content (TCC) is reported as the mean percentage of inflorescence dry mass ± SE.

| ADMIXTURE<br>cluster<br>location | genotype summary |  |  | cannabinoid content |  |  |  |
| --- | --- | --- | --- | --- | --- | --- | --- |
|  | *N<sub>g</sub>* | *f* | *CBDAS* | *N<sub>c</sub>* | %THC (SE) | %CBD (SE) | %TCC (SE) |
| **Mississippi R (1)** |  |  |  |  |  |  |  |
| WI, Cross Plains | 22 | 0.48 | $C^F C^F$ | 4 | 0.04 (0.00) | 0.58 (0.15) | 0.72 (0.16) |
| T7N R7E | 13 | 0.28 | $C^F C^X$ | 16 | 0.13 (0.02) | 0.36 (0.05) | 0.66 (0.08) |
| | 11 | 0.24 | $C^X C^X$ | 8 | 0.25 (0.03) | 0.02 (0.00) | 0.51 (0.06) |
| WI, West Point | 6 | 0.15 | $C^F C^F$ | 2 | 0.10 (0.05) | 0.59 (0.02) | 0.87 (0.15) |
| T10N R7E | 24 | 0.59 | $C^F C^X$ | 24 | 0.25 (0.02) | 0.74 (0.05) | 1.31 (0.08) |
| | 11 | 0.27 | $C^X C^X$ | 10 | 0.37 (0.03) | 0.04 (0.00) | 1.05 (0.09) |
| **WNC-a (2)** |  |  |  |  |  |  |  |
| ND, Waldo | 44 | 0.88 | $C^F C^F$ | 7 | 0.07 (0.01) | 1.72 (0.21) | 1.94 (0.23) |
| T130N R49W | 4 | 0.08 | $C^F C^X$ | 6 | 1.16 (0.07) | 1.63 (0.14) | 3.19 (0.17) |
| | 2 | 0.04 | $C^X C^X$ | - | - | - | - |
| SD, Cavour | 8 | 0.80 | C<sup>F</sup>C<sup>F</sup> | 5 | 0.08 (0.01) | 2.18 (0.29) | 2.47 (0.32) |
| T111N R60W | 2 | 0.20 | C<sup>F</sup>C<sup>X</sup> | 2 | 1.03 (0.43) | 1.40 (0.58) | 2.78 (1.13) |
|  | - | - | C<sup>X</sup>C<sup>X</sup> | - | - | - | - |
| **WNC-b (3)** |  |  |  |  |  |  |  |
| IA, Remsen | 7 | 0.70 | C<sup>F</sup>C<sup>F</sup> | 7 | 0.10 (0.01) | 2.17 (0.30) | 2.44 (0.33) |
| T92N R43W | 3 | 0.30 | C<sup>F</sup>C<sup>X</sup> | 3 | 0.47 (0.06) | 0.78 (0.33) | 1.46 (0.43) |
|  | - | - | C<sup>X</sup>C<sup>X</sup> | - | - | - | - |
| MO, Clark | 9 | 0.75 | C<sup>F</sup>C<sup>F</sup> | 4 | 0.09 (0.01) | 1.14 (0.25) | 1.40 (0.30) |
| T64N R40W | 2 | 0.17 | C<sup>F</sup>C<sup>X</sup> | 3 | 0.59 (0.24) | 0.85 (0.22) | 1.67 (0.49) |
|  | 1 | 0.08 | C<sup>X</sup>C<sup>X</sup> | - | - | - | - |
| **Indiana (4)** |  |  |  |  |  |  |  |
| IN, Wayne | 12 | 0.24 | C<sup>F</sup>C<sup>F</sup> | 11 | 0.06 (0.01) | 0.94 (0.15) | 1.12 (0.16) |
| T32N R3W | 21 | 0.43 | C<sup>F</sup>C<sup>X</sup> | 28 | 0.40 (0.02) | 0.48 (0.02) | 1.02 (0.04) |
|  | 16 | 0.33 | C<sup>X</sup>C<sup>X</sup> | 10 | 0.72 (0.07) | 0.02 (0.00) | 0.90 (0.08) |
| IN, Barkley | 5 | 0.20 | C<sup>F</sup>C<sup>F</sup> | 5 | 0.07 (0.01) | 0.99 (0.15) | 1.17 (0.17) |
| T30N R6W | 10 | 0.40 | C<sup>F</sup>C<sup>X</sup> | 14 | 0.49 (0.06) | 0.57 (0.06) | 1.19 (0.12) |
|  | 10 | 0.40 | C<sup>X</sup>C<sup>X</sup> | 6 | 0.86 (0.14) | 0.03 (0.01) | 1.04 (0.16) |

### Linkage Disequilibrium

LD decay occurred rapidly with increasing physical distance. The average maximum *r*^2^ value was 0.47. As LD decayed to its half maximum (*r*^2^ = 0.24), the corresponding physical distance was approximately 6.7 kb (Supplementary Figure 5).

### *CBDAS* Genotyping

Among US feral *Cannabis* collections, *CBDAS* genotypes were resolved for approximately 1,400 accessions from 90 populations located among 12 states into one of three diploid classes (Supplementary Table 6). Among all accession genotypes, 1120 were homozygous for functional (C^F^C^F^), 91 were homozygous for non-functional (C^X^C^X^), and 208 were heterozygous (C^F^C^X^) comprising study wide allele frequencies of 0.86 and 0.14 for functional (C^F^) and non-functional (C^X^) *CBDAS*, respectively (Supplementary Table 7). Among a subset of 53 populations with >= 10 resolved genotypes, functional *CBDAS* (C^F^) allele frequency ranged from 0.99 to as low as 0.4 among 36 populations and was fixed at 1.0 among 17 populations (Supplementary Table 7). The *CBDAS* allele frequency among this population subset varied geographically across a region comprising 10 contiguous upper midwestern states (Figure 5). Populations from which accession genotypes were exclusively homozygous for functional *CBDAS* were located in all but two states, North Dakota and Indiana, at the northeastern and southeastern margins of the upper Midwest sampling region. A total of five populations with exclusively functional *CBDAS* alleles were also composed of individuals belonging entirely to the Mississippi River group. In addition, three populations outside of the upper Midwest, two in Colorado and one in New York, were completely homozygous for functional *CBDAS* (Supplementary Table 8).

**Figure 5.**
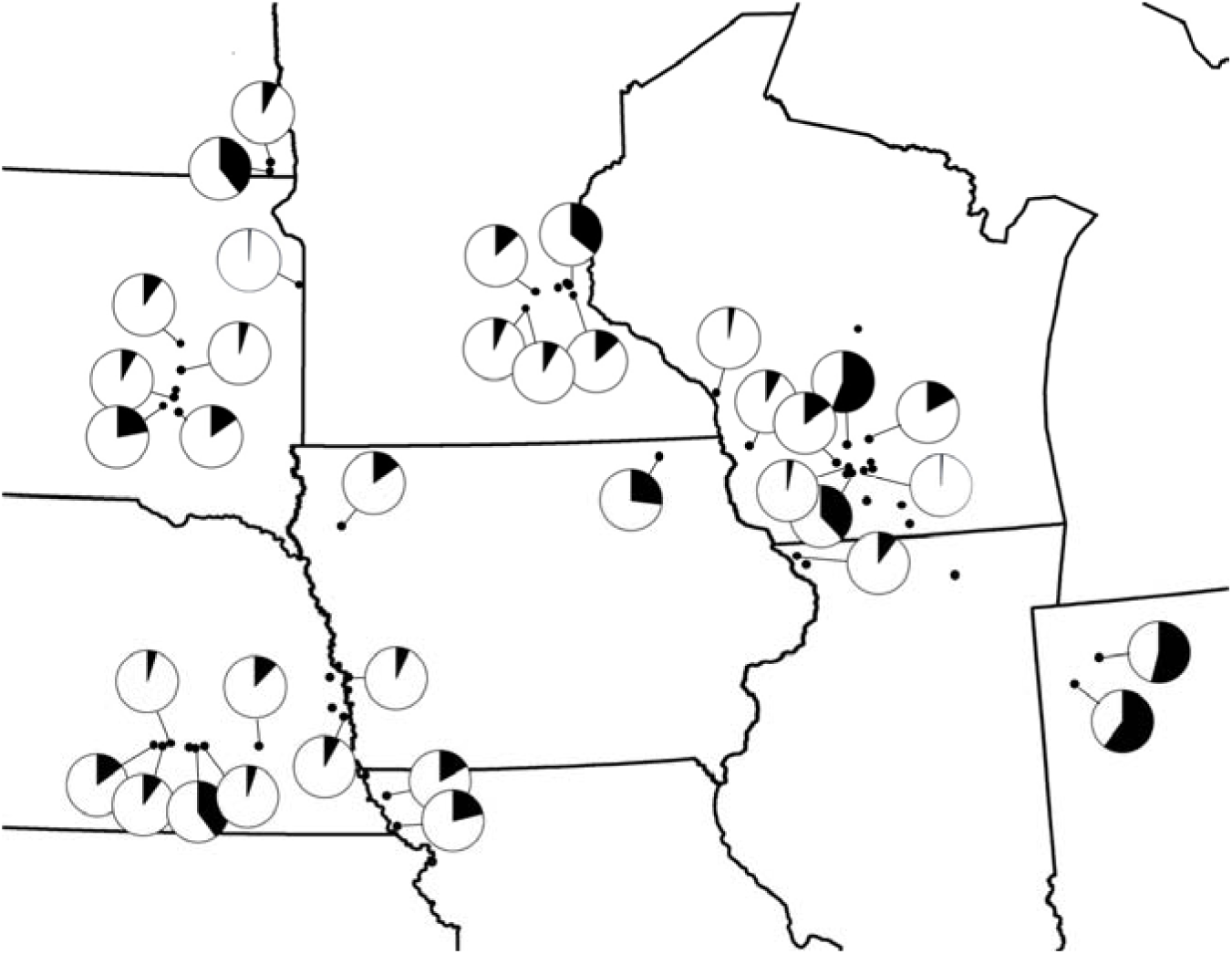
*CBDAS* allele frequency variation among populations with >/= 10 resolved genotypes. Population localities are indicated by small black circles. Pie-charts display the proportion of functional (C^F^; white) and non-functional (C^X^; black) *CBDAS* alleles observed within populations among individuals for which a *CBDAS* genotype was resolved. Populations not linked to a chart are fixed where *f*C^F^ = 1.

### Cannabinoid Analysis

Among US feral *Cannabis* collections across the 10 states comprising the upper Midwest sampling region (Figure 1b), female floral tissues (pistillate inflorescences) from 523 accessions were analyzed by gas chromatography for determination of the % dry weight content of eight cannabinoid compounds except for 30 accessions for which determinations included seven compounds (omitting Δ8-THC; Supplementary Table 6). When categorized by the ratio of Δ9-THC:CBD content, these accessions included 91 THC-type, 233 intermediate-type, and 244 CBD-type feral *Cannabis* plants selected from 61 populations (Supplementary Table 8). Total cannabinoid content ranged 0.23-2.96%, 0.29-4.09%, and 0.21-4.73% among accessions classified as THC-type, intermediate-type, and CBD-type, averaging (±SE) 0.98% (0.07), 1.38%, and 1.50%, respectively (Supplementary Table 8). Histograms of total cannabinoid content among 523 feral *Cannabis* accessions are shown in Figure 6, along with the histograms for a subset of accessions meeting criteria for assignment to one of five ADMIXTURE groups.

**Figure 6.**
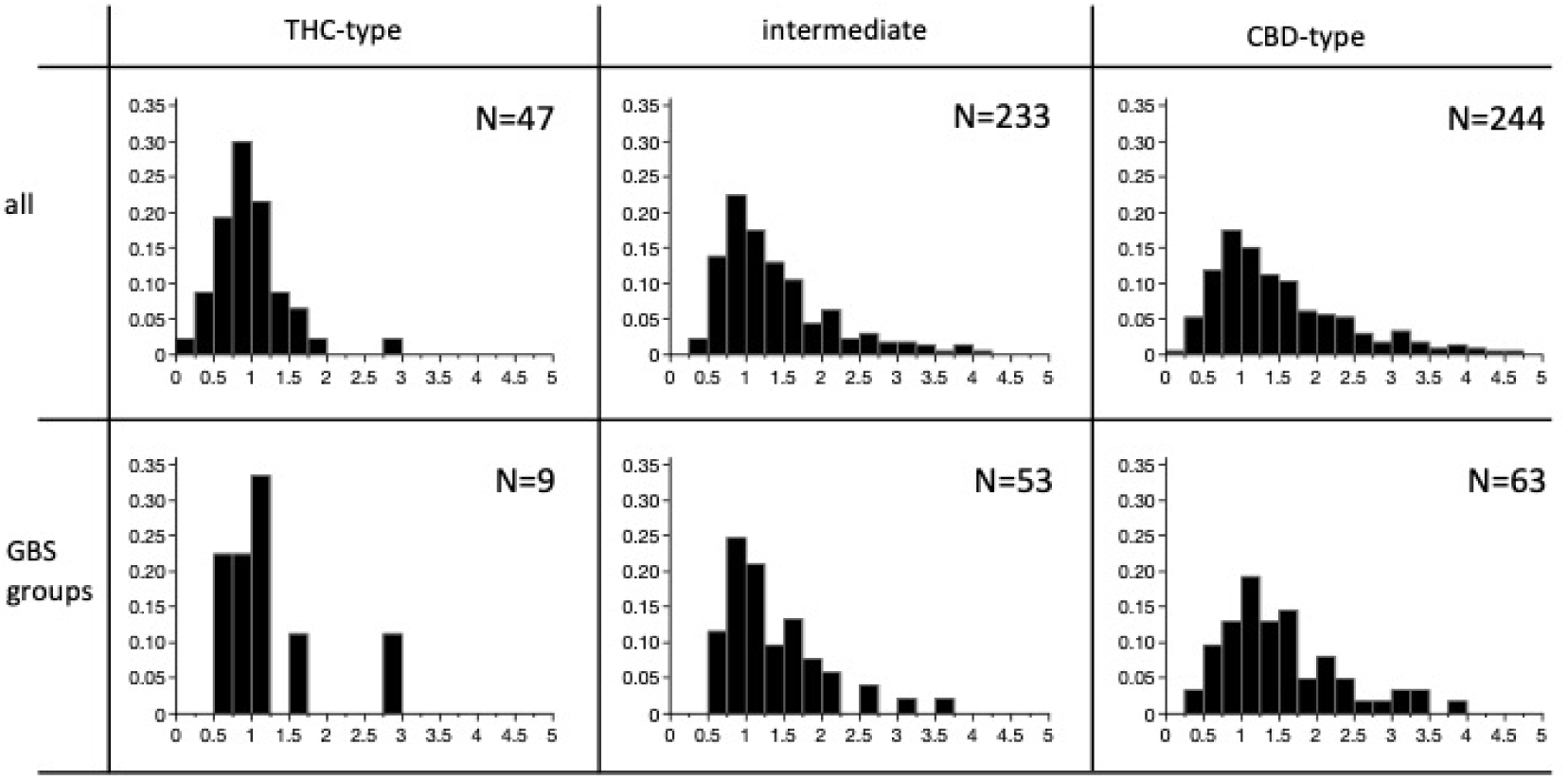
Distribution of total cannabinoid content (% dry weight of “female flower” or pistillate inflorescence) among plants differing in the ratio of THC:CBD. Left, center, and right columns include plants with either an excess of THC (THC-type or type-I), similar amounts of THC and CBD (intermediate or type-II), or an excess of CBD (CBD-type or type-III), respectively. Upper row histograms include all accessions for which cannabinoid content was measured by gas chromatography where N indicates the total number of plants. Lower row histograms include a subset of genotyped plants assigned to each of the three types using the PACE assay. Histograms are proportional to the number of samples sharing ±0.5% cannabinoid content.

## DISCUSSION

US hemp breeding programs have been legislatively impeded for nearly a century due to federal prohibition under the 1937 Marihuana Tax Act and the Controlled Substances Act of 1970. Federal drug law policies prohibited the cultivation and use of *Cannabis* in any form, crippling potential industrial and medicinal use. Interests in industrial hemp production increased with the passage and expansion of the 2014^12^ and 2018 Farm Bill^13^. Still, the limited availability of complaint and diverse *Cannabis* germplasm represents a major bottleneck for breeding and research efforts.

Although the last legal US hemp crops were planted in 1958^37,38^, remnants of domesticated industrial hemp have escaped to form naturalized, thriving feral populations across regions of the US where it had been previously cultivated. Additional dispersal of seed by birds, animals, and anthropogenic activities might have resulted in further establishment in regions where hemp had not previously been cultivated. Sites suggestive of such secondary range expansion include roadsides, railroad rights-of-way, livestock feedlots, and agricultural boundaries especially as associated with trees and utility lines. We relied on university collaborators, farmers, and citizen scientists to identify and/or collect feral *Cannabis* plants across the US based on sites recorded in iNaturalist or received by word of mouth. Whereas site identification was essentially straightforward, collection of samples was occasionally challenging because of residual stigma associated with cannabis. Additionally, occurrences within interstate highway corridors, on private property, and accompanied by dense vegetation made collection difficult. Wider identification of feral *Cannabis* occurrence and collection is particularly needed in the West, South, and Northeast geographical regions of the US. Meeting this need will require effective communication about the potential economic benefits of adopting industrial hemp production among existing agricultural crops.

Five subpopulations were identified among US feral *Cannabis* collections. Generally, samples clustered according to state of origin showing that geographical location explains genetic variation. Most notable were two clusters, one containing accessions primarily from Indiana and the other containing accessions from New York. The remaining three clusters were less variable and contained samples from states along the Mississippi River basin and within West North Central states.

Pairwise comparison of fixation indices (F_ST)_ between subpopulations found genetic distance to be in accordance with geographical location with the most divergent subpopulations from Indiana (group 5) and New York (group 3). The F_ST_ values from this study are higher than those previously recorded in numerous studies focused on drug-type *Cannabis*^26,27,29,34^, which is likely attributed the broader genetic background of hemp as compared to drug-type *Cannabis*. The least divergent subpopulations (F_ST_ = 0.07 - 0.09) may have resulted from a single introgression to the Upper Midwest during the WWII “*hemp for victory”* era followed by dispersal by humans or animals across the sampled regions. Interestingly, many of the collection sites in these regions were either along railroad rights-of-way or livestock facilities. High F_ST_ between the Indiana and New York groups indicate separate introgressions. Indeed, hemp arrived at different geographical regions at distinct periods throughout history with samples arriving in the Northeast as early as the 16^th^ century, Kentucky in the 18^th^ century and Wisconsin during the 20^th^ century^19,39,40^. The three primary subpopulations identified in the current study may have originated from distinct introgression and feralization events and therefore harbor unique alleles based on origin.

High pairwise F_ST_ scores were found for all comparisons indicating regions of the genome that may be under selective pressures across different regions. The highest F_ST_ values were associated with genomic regions on chromosomes 4 and X (Figure 4; Supplementary Figure 4). Previous work identifies similar peaks and hypothesizes that the regions may be the result of chromosomal inversions maintained at high frequency in the feral US populations^34^. The *Cannabis* pangenome has confirmed a large-scale inversion on chromosome 4 in a feral collection from Boone County, Iowa as compared to high cannabinoid cultivars^41^. We find this same elevated F_ST_ primarily in samples from the West North Central-a and Indiana groups suggesting it is segregating within US feral populations. Low global F_ST_ observed between West North Central-a and -b groups, indicates these accessions are mostly differentiated by this inversion. Analysis of the 15 Mb region surrounding the chromosome 4 inversion explains 65% of the variation among samples in the PCA (Supplementary Figure 4). Because this inversion is found in some frequency across all subpopulations, it is likely this inversion occurred prior to the introgression of hemp to the United States. Fixation in several populations suggests that it may confer a selective advantage in certain environments.

The expected heterozygosity level (0.20) for the accessions in our study was similar to the observed heterozygosity (0.20) indicating the populations are not experiencing significant selective forces such as inbreeding or genetic draft. Our observed heterozygosity is more similar to what has been previously described in hemp (0.27-0.16) as compared to drug-type cannabis (0.12-0.14)^26,28^. A lower observed heterozygosity was observed in the New York subpopulation which may be attributed to its small population size and geographic isolation. The negative value of inbreeding coefficients recorded in the MS-River, WNC-a and WNC-b subpopulations, imply increased heterozygosity and higher genetic diversity when compared to the Indiana and New York feral values reflecting random mating and genetic stability among the populations. Polymorphism Information Content (PIC) is a measurement of how well a genetic marker can distinguish different individuals in a population. It is a key indicator of marker quality in genetic studies and is used to predict the likelihood of a marker’s heterozygous genotype being passed on to offspring^42^. The PIC marker values per population ranged between 0.15 – 0.30, indicating they are very informative at discriminatory analysis between feral populations.

High genetic variability from AMOVA as observed within accessions followed by between accessions within groups and lowest between groups suggests potential for trait discovery in breeding programs. The moderate differentiation between groups reiterates the high heterozygosity often recorded in *Cannabis* associated with wind pollination. Within accession AMOVA values among US feral collections was higher than that of the Spanish and Moroccan *Cannabis* germplasm studies^34,43^. Discrepancies in variation contributed by these studies are consistent with other studies and may be due to differences in sample size, sample composition and types of markers used.

Linkage disequilibrium decayed rapidly (6.7kb) in the feral germplasm. This was similar to a recent report using a mixture of hemp and drug-type ferals (LD = 6.0 kb)^29^ and faster than in a recent study of drug-type *Cannabis* averaging between 22.6 to 89.0 kb across chromosomes^44^. Historic breeding efforts during the 18^th^ and 19^th^ centuries that entailed intentional hybridization as well as wind dispersal of hemp pollen over great distances may account for the plunge in LD decay of feral *Cannabis* plants. This finding will be important for finding marker-trait associations and developing genotyping resources to adequately cover haplotype blocks.

Total cannabinoid content in Midwestern feral *Cannabis* was rather low compared to drug cultivars although somewhat higher than hemp cultivars^45^ with population averages (±SE) ranging from 0.62 (0.05) to 4.04 (0.26) (Supplementary Table 8). Most populations contained at least some individual plants with THC content above the US statutory threshold of 0.3%. Such plants were either of intermediate (C^F^C^X^) or THC-type (C^X^C^X^) genotypes as indicated by the presence of non-functional *CBDAS* alleles at low frequency. Approximately one third of 53 populations with at least 10 resolved accession genotypes were fixed for functional *CBDAS*, with mixed populations varying in non-functional allele frequency from >0.1 to 0.6. These observations are consistent with evidence from Minnesota^35^ suggesting that midwestern “feral hemp” does not neatly fit the current statutory definition of hemp but is more accurately described as feral *Cannabis*. Genotype frequencies in mixed populations were generally skewed in favor of CBD-type, with a lower frequency of intermediates and the occasional THC-type plant. There is little reason to expect that the original *Cannabis* introductions to the region in the 19th century for fiber production were pure hemp, considering that the chemical nature of intoxicating versus non-intoxicating *Cannabis* (THC versus CBD) was not discovered until decades later^46,47^. Populations at or near Hardy-Weinberg equilibrium with relatively low yet varying non-functional *CBDAS* allele frequencies might be interpreted as subject to local genetic drift with little or no selection on cannabinoid content and/or little immigration of THC-type genetics from elsewhere. However, several populations deviated from this pattern with a high proportion of intermediate-type plants or a non-equilibrium excess of THC-type plants, which might be interpreted as the result of introgression from more recent drug-type cultivation. Genotyping of historical specimens from herbarium collections might shed light on how and whether cannabinoid genetics has changed over time^48^.

*CBDAS* genotypes as measured by the PACE assay^49^ accurately predicted each THC:CBD ratio class in most cases (87%). In three cases, cannabinoid content was so close to the limit of detection by gas chromatography (0.01%) that the THC:CBD ratio lacked meaning, and we do not consider them to challenge the accuracy of the genetic assay. However, 33 of 465 accessions (13%) had *CBDAS* genotypes that did not correlate with the THC:CBD ratio even after independent, repeated measures. Such plants merit further investigation of the possibility that allelic variation at the primer binding sites could skew the assay. This seems more likely than a departure from the simple genetic mechanism of codominant inheritance^11^ as others have suggested^50^. Another limitation of the PACE assay that we observed when applied to feral *Cannabis* was a relatively high failure rate (20%) in which some accessions failed to amplify (18%) while others (2%) yielded ambiguous fluorescence patterns where it was not possible to distinguish between the intermediate genotype and a homozygous state.

Feral accessions are important genetic reservoirs of useful alleles and superior traits for the improvement of *Cannabis* plants for fiber, grain or cannabinoids^51,52^. The persistence of these feral *Cannabis* populations in selected habitations and even spread to new localities after decades of naturalization attest to their resilience and importance as new tools for breeding improved cultivars with increased adaptation to biotic and abiotic stress factors. The conservation of these *Cannabis* collections in the USDA-ARS Hemp Germplasm Repository will ensure the long-term security of these genetic resources against any future political or societal pressures and their genetic integrity will be continually maintained through thorough regeneration practices. Conservation of these diverse feral *Cannabis* materials in the genebank promises in contrast to historic absence under decades-long prohibition, to provide access to diverse and compliant *Cannabis* germplasm.

## MATERIALS AND METHODS

### Germplasm collection

To identify feral *Cannabis* sites, we used the iNaturalist application (a joint initiative by the California Academy of Sciences and the National Geographic Society), an American nonprofit social network of citizen scientists and biologists who identify and share information about biodiversity across the globe. Additional locations were shared by collaborators via word of mouth. Seeds and floral tissue from 1,815 female hemp plants from 88 unique populations were collected across twelve states in 2022 and 2023 (Figure 1a; Supplementary Table S1; Supplementary Material). An additional feral population from New York was obtained from USDA-ARS (Geneva, NY) and seeds from two Colorado populations were contributed from Colorado State University.

### Sample preparation and genotyping

A subset of 760 samples, representing twelve states and ecogeographic US locations, were selected for genotyping. Leaf tissue from individual plants was lyophilized (Labconco, Kansas City, MO, USA), then approximately 20 mg dry weight was homogenized (SPEX SamplePrep, Metuchen, NJ, USA), and genomic DNA was extracted using NucleoSpin 96 Plant II core kit (Macherey-Nagel GmbH & Co., Duren, Germany). DNA quality was checked by 1% agarose electrophoresis and quantified with NanoDrop ND-1000 Spectrophotometer (Thermo Fisher Scientific, USA). The feral hemp DNA quantification, library preparation, and sequencing was constructed using the genotyping-by-sequencing (GBS) protocol^53^ at the University of Wisconsin-Madison Biotechnology Center. In summary, Illumina adapters and sample-specific barcodes were annealed after restriction enzyme digestion using *ApeKI*. Multiplexed samples were then sequenced on Illumina NovaSeq 6000 (Illumina, Inc., San Diego, CA, USA), generating on average 3.5 million, 150 bp paired end reads per sample.

### Variant discovery and data filtering

The Tassel GBS Discovery Pipeline v2^54^ was used for variant detection, using the cs10 reference genome v2^9^ (Genbank assembly accession ID = GCF_900626175.2) with BWA-MEM^55^. Raw VCF files were filtered for SNPs > 10 % missing data, < 5 % minor allele frequency, > 10 % missing individuals, and finally pruned to reduce redundancy by removing SNPs that are highly correlated with others (--geno 0.1, --maf 0.05, --mind −0.1, --indep-pairwise 50 2 0.2) using PLINK V.1.90b6.21^56^. The final VCF file contained 21,037 SNPs (∼ 21K) used in downstream analyses. Chromosome-wide SNP count and density plots were performed (https://github.com/YinLiLin/R-CMplot) in the R environment (version 4.3.2).

### Population structure and principal component analysis

Population structure was determined using ADMIXTURE^57^ to estimate the genetic ancestries of each feral accession, and cross-validation plots were observed for the optimum number of clusters. ADMIXTURE identifies *K* genetic clusters, where *K* is specified by the user (K2-K40), from the provided SNP data. For each individual, the ADMIXTURE method estimates the probability of membership to each cluster using Bayesian Information Criterion. Low admixture samples were assigned by setting the membership coefficient to 0.7. Eigenvalues and eigenvectors for genome-wide principal component analysis (PCA) were generated using PLINK open-source analysis toolset and then plotted in the R-software^58^, package “ggplot^59^.

### Genetic differentiation and diversity indices

To measure genome-wide genetic differentiation (F_ST_) and molecular variance (AMOVA) among identified subgroups, poppr^60^ and *Hierfstat*,^61^ R packages were employed. Diversity indices (*Ho* = Observed heterozygosity, *He* = Expected heterozygosity, MAF = Minor allele frequency, PIC = Polymorphic information content, and F_IS_ = Inbreeding coefficient) were estimated with PLINK. To estimate localized genetic differentiation across the genome, pairwise F_ST_ was calculated using VCFtools with a 100,000 bp, non-overlapping sliding window and plotted in R using package “ggplot”.

### Linkage disequilibrium

The LD squared allele frequency correlation (r^2^) estimate for each pairwise SNP comparison was generated in TASSEL 5.0 and visualized by plotting r^2^ values against physical distance in R package “ggplot”. A nonlinear regression curve was used in the estimation of LD decay. The LD decay rate was determined by the point of intersection with the LD curve and half its maximum value (*r*^2^ = 0.24).

### *CBDAS* locus genotyping

To resolve *CBDAS* locus genotypes of accessions, we used the CCP1 primer set^49^ that has been validated to detect and distinguish presence of functional (C^F^) and non-functional (C^X^) psuedogenous alleles of *CBDAS* that are tightly and opposingly linked to non-functional pseudogenes and a functional allele of the *THCAS* gene^9,11^. PACE™ 2.0 2X master mix with standard ROX was used according to the manufacturer’s protocol (3CR Bioscience Limited) to produce fluorescently labeled *CBDAS* locus real-time PCR fragments on Bio-Rad CFX96, Roche 980, or Thermo-Fisher QuantStudio qPCR instruments following respective PACE^TM^ protocols for bi-allelic genotyping. Genomic DNA from feral *Cannabis* accessions were prepared as assay templates from dried leaf or floral tissues using either Extract-N-Amp^TM^ Plant PCR (Millipore Sigma) extraction reagents or using a Nucleospin 96 Plant II core kit (Macherey-Nagel GmbH & Co., Duren, Germany) kit according to the respective manufacturer’s protocol.

### Chemical cannabinoid diversity

Dried floral tissue samples were analyzed using gas chromatography^62^ to measure the percentage of total inflorescence dry mass of: cannabichromene (CBC), CBD, cannabigerol (CBG), cannabinol (CBN), delta-8-tetrahydrocannabinol (d8-THC), THC, and tetrahydrocannabivarin (THCV). We report total cannabinoid content as the sum of these individual compounds as well as the percent CBD and THC content. Cannabinoid profiling was conducted on up to five plants of each genotype, as determined by the PACE assay, per population.

## Supporting information

Supplementary word document

## Acknowledgements

The authors express their sincere appreciation to Joseph Smeenge and Pawan Basnet for assistance with fieldwork and genotyping, citizen scientist contributors to iNaturalist, the S1084 Multistate Research Project and every individual that helped to identify potential collection sites or collected and sent feral *Cannabis* seeds and/or leaf tissue. We also thank the United States Department of Agriculture – National Institute of Food and Agriculture (USDA-NIFA), Supplemental and Alternative Crops (SAC) grants program (2021-38624-35738) and USDA-ARS agreement number 58-5090-2-03 for funding this research. This work was also supported in part by awards 3000032227 and 3000038264 from the Agricultural Growth, Research, and Innovation (AGRI) Program of the Minnesota Department of Agriculture. Lastly, Weiblen is grateful to the offices of the General Counsel and the Vice President for Research at the University of Minnesota, and the U. S. Drug Enforcement Administration for guidance in developing a field protocol for feral *Cannabis* collection.

## Author contributions

SE and GW conceived the research plan. AA, JW, ES, CM, ME, GW and SE performed the research and analyzed the data. AA, SE, JW, and GW wrote the manuscript with input from all the authors. All authors edited and approved the final version of the manuscript.

## Data availability statement

DNA sequencing data are available through NCBI’s BioProject ID: PRJNA1206134. All other data are available upon request or provided in the supplementary material.

## Declaration of competing interest

The authors declare that they have no known competing financial interests or personal relationships that could have appeared to influence the work reported in this paper.

## SUPPLEMENTARY FIGURES

**Supplementary Figure 1:**
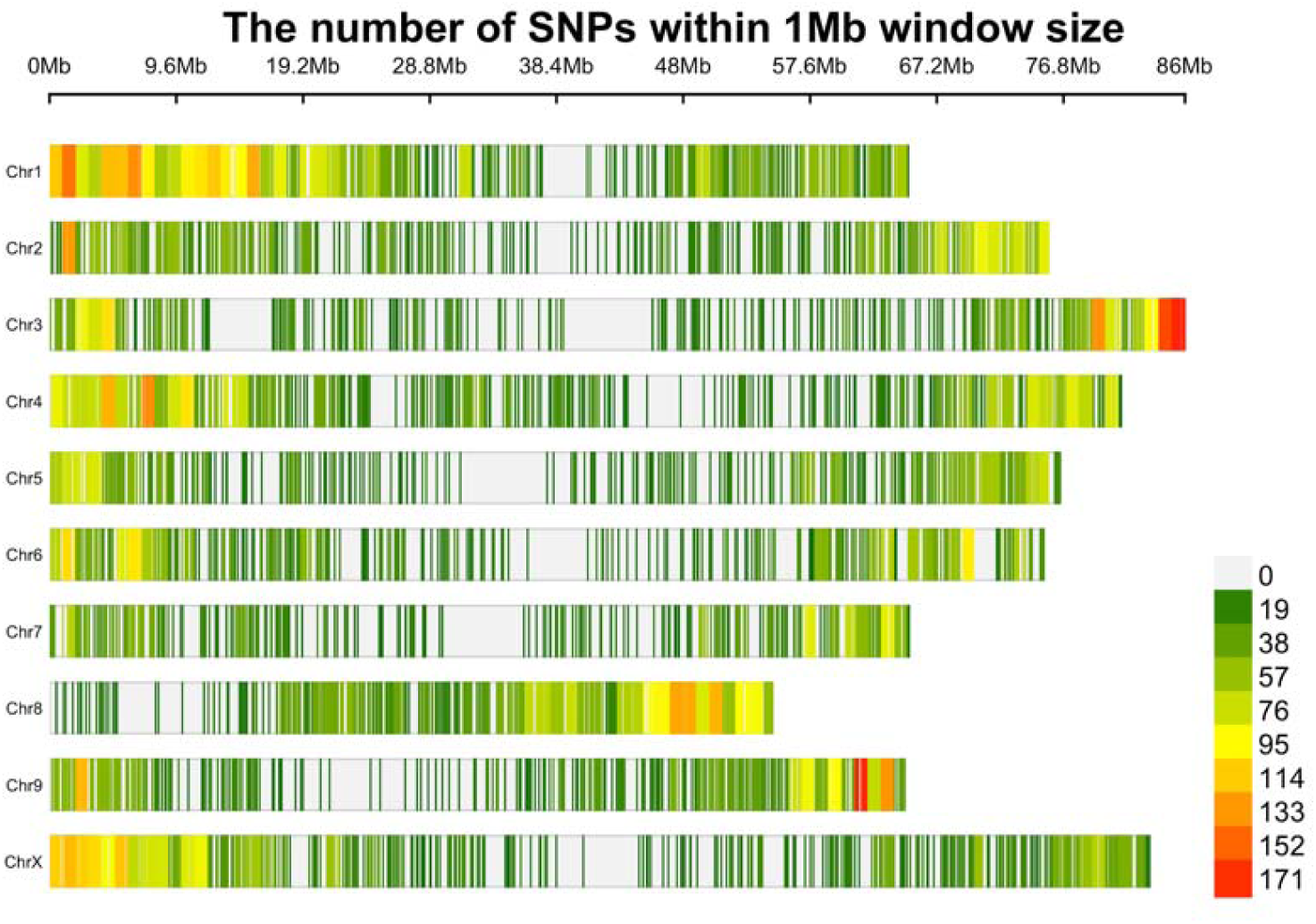
SNP density plot of the chromosome-wise number of SNPs within 1 MB window size.

**Supplementary Figure 2:**
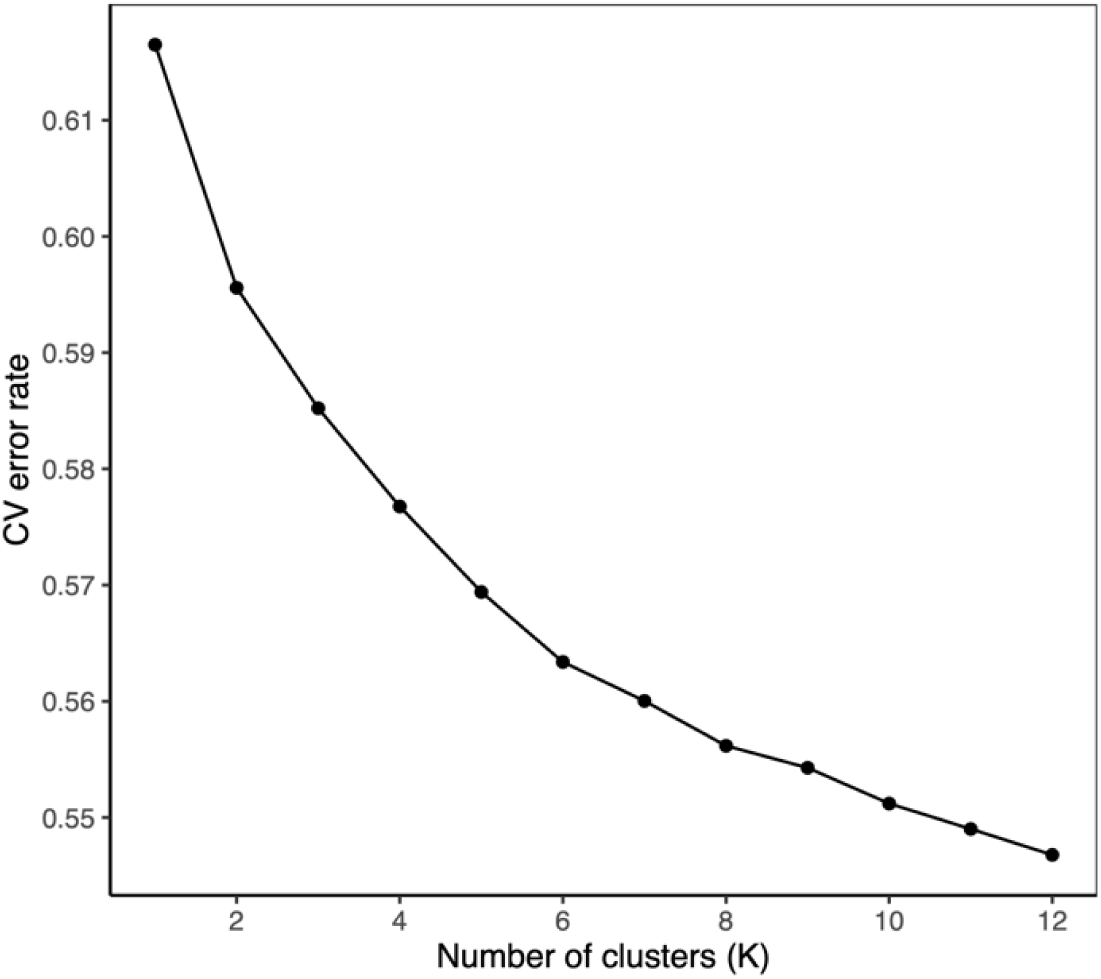
Plot of cross-validation error against number of clusters (K)

**Supplementary Figure 3:**
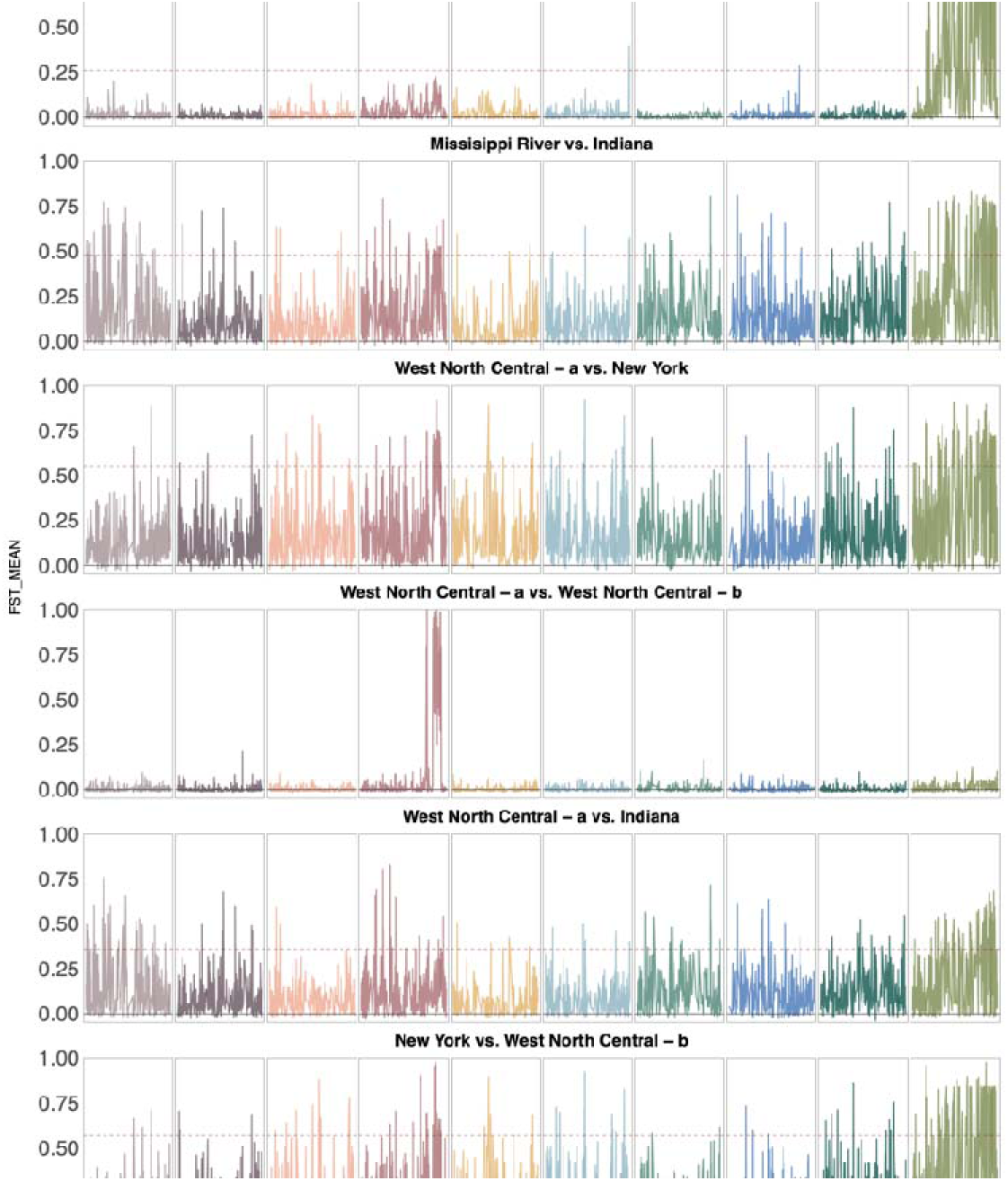
Sliding window FST for all pairwise comparisons of the 5 subgroups.

**Supplementary Figure 4:**
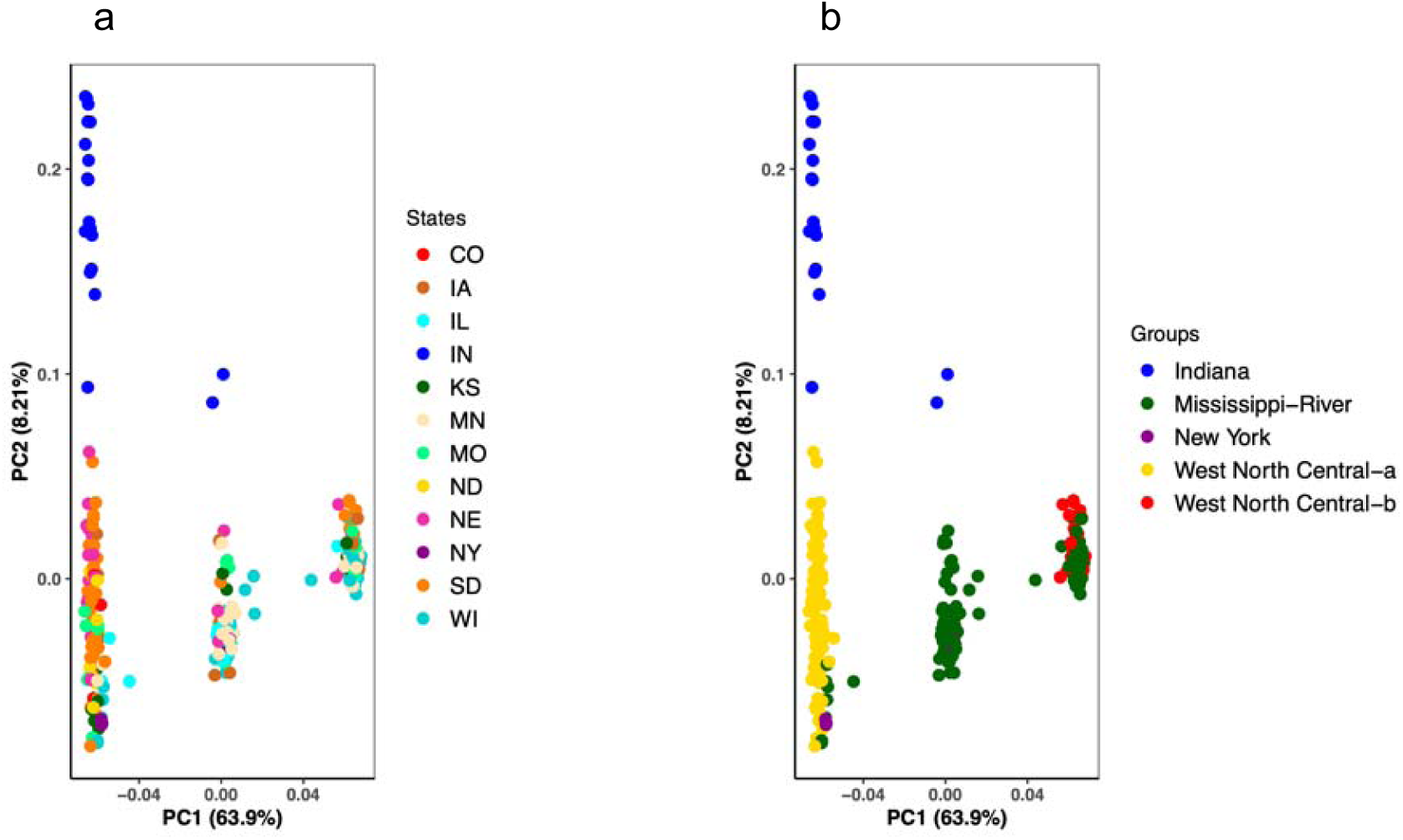
PCA for SNPs within 60-95 Mb region on chromosome 4 colored by a) states and b) ADMIXTURE groups.

**Supplementary Figure 5:**
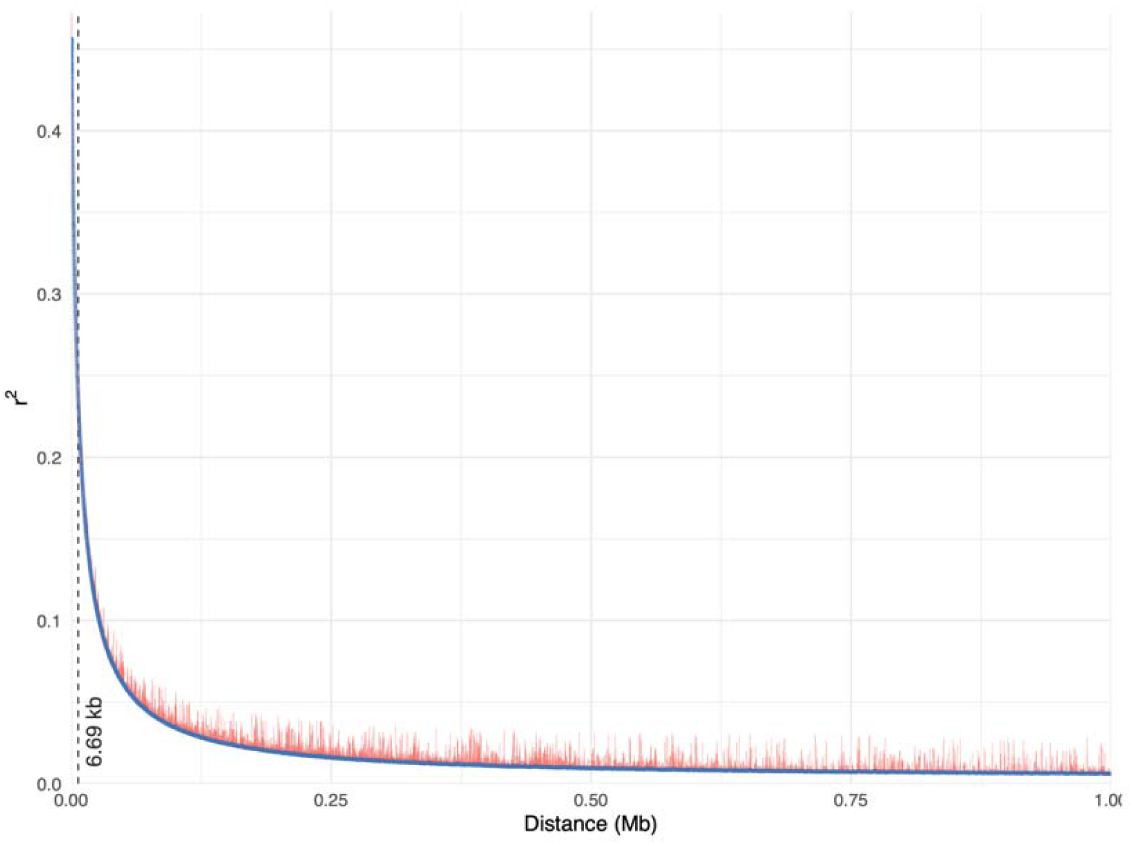
Genome-wide linkage disequilibrium (LD decay) plot. The *X*-axis indicates distances, and the *Y*-axis indicates pairwise correlations (r^2^) between all pairs of markers

## Notes

### Competing Interest Statement

The authors have declared no competing interest.

