## Supplementary word document for "Genetic Diversity, Population Structure, and Cannabinoid Variation in Feral *Cannabis sativa* Germplasm from the United States"

**Feral Cannabis Collection Protocol**

**Materials needed:**

Grocery paper bags #10 (6.5 x 4 x 13.25) (50 per population)

Reflective vests

Waterproof folders/bags for datasheets

Datasheets

Sharpies

Pens

**Tools required:**

Gloves

Tape measure

Camera (Phone’s camera is okay)

Loppers

Pruning shears

**Feral Cannabis Collection Protocol:**

1. Secure state or federal licensing to possess industrial hemp in your jurisdiction. ***UW Madison has a federal license and will provide it to collectors***
2. Maintain copies of permits, licenses, and/or written authorization during fieldwork and with collected material through all subsequent steps in the protocol.
3. Investigate locations identified by citizen scientists (either through iNaturalist or word of mouth) as potential feral hemp populations. Avoid cultivated fields of industrial hemp (with potential proprietary genetics) and cultivated marijuana, be it state-registered or clandestine and illegal. Endeavor to cover the broadest range of ecoregions that can feasibly be accessed during fieldwork.
4. Secure verbal and/or written permission from the landowner/tenants to collect plant material.
5. Collect GPS coordinates and take pictures of the plants while capturing their immediate surroundings using a smartphone. Upload coordinates along with pictures on iNaturalist for future collection purposes. Record a brief description of the site – drainage patterns, surrounding vegetation, elevation, and soil classification from the web soil survey on the provided datasheet.
a. Preferred population size >50 female individuals, at a minimum of 10 females.
b. Populations separated by at least 5 miles (8km) are preferred to increase the likelihood of pollen isolation.
c. Give each population an ID code in this scheme: state (WI), year (2), collector’s initials (SE), two-digit population number (01), and plant number (01) -i.e. WI-23-SE-01-01.
d. Upload photos to a safe drive for backup
6. Locate up to 50 female (seed-bearing) plants. For each plant, label a large paper bag with an ID code. Chop down at the base nearest to the ground with the loppers, lay on the ground, and measure the height with a tape measure, record the ID code and height in the nearest inch on the datasheet. Use the pruning shears to cut the terminal and lateral seed-bearing branches into 6-8” sections and fill the bag up to ¾ of its height, allowing enough space to fold the top of the bag over. Do not collect more material than would fit a single bag. Collecting each plant in a single, separate bag is essential. If there are more seed-bearing branches on a plant than fit the bag, you might prioritize the most heavily seeded branches but do not overstuff the bag and do not put material from multiple plants in the same bag.
7. Hot, moist conditions are detrimental to seed viability. Bagged material should begin drying within 12 hours of collection. At no point should the paper bags be enclosed in plastic. Samples should be kept as cool as possible during transport from the field to a location where they can be dried.
8. Dry the bags at room temperature with a fan to move air for at least one week. Seed and plant material can be placed in a drying oven as long as temperatures are less than 95°F. Conditions exceeding 95°F are detrimental to viability.
9. Once the plants are dried, hand-strip the branches to remove the bulkier stems from each bag. Stems may be discarded in compost or as waste. Fold the top of the bag over and staple or tape it closed.
10. Line in a shipping box with a plastic bag, place paper bags and the datasheet(s) inside the plastic bag, seal the box. Attach the shipping label provided and arrange for a Fedex pickup or deliver to a Fedex drop-off location. Refer to the contents only as “Dried plant material for scientific research. Not hazardous and of no commercial value.”

**Data collection sheet should have the following details:**

Collector’s name; Date; Location; Latitude; Longitude; State; Population #; Site Description

Column Headers: Plant, Height (specify cm or inches)

01, 159

02, 120

03, 114

04, etc
